# Arabidopsis HB52 mediates the crosstalk between ethylene and auxin signaling pathways by regulating *PIN2, WAG1, and WAG2* during primary root elongation

**DOI:** 10.1101/246017

**Authors:** Zi-Qing Miao, Ping-Xia Zhao, Jie-Li Mao, Lin-Hui Yu, Yang Yuan, Hui Tang, Cheng-Bin Xiang

## Abstract

The gaseous hormone ethylene participates in many physiological processes of plants. It is well known that ethylene-inhibited root elongation involves basipetal auxin delivery requiring PIN2. However, the molecular mechanism how ethylene regulates *PIN2* is not well understood. Here, we report that the ethylene-responsive HD-Zip gene *HB52* is involved in ethylene-mediated inhibition of primary root elongation. Using biochemical and genetic analyses, we demonstrated that *HB52* is ethylene-responsive and acts immediately downstream of EIN3. *HB52* knock-down mutants are insensitive to ethylene in primary root elongation while the overexpression lines have dramatically shortened roots like ethylene treated plants. Moreover, HB52 upregulates *PIN2, WAG1*, and *WAG2* by directly binding to their promoter, leading to an enhanced basipetal auxin delivery to the elongation zone and thus inhibiting root growth. Our work uncovers HB52 as an important crosstalk node between ethylene signaling and auxin transport in root elongation.

## Introduction

Ethylene is a gaseous phytohormone which regulates a multitude of processes at trace levels. It is well known for triggering the shedding of leaves, the ripening of fruits, and the defense of plants. It also plays an indispensable role in root development (Alonso and Ecker, 1999; Grbić and Bleecker, 2003; Chaves and Mello-Farias, 2006; Ruzicka et al., 2007). Exogenous treatment with ethylene (C_2_H_4_) or its biosynthesis precursor 1-aminocyclopropane-1-carboxylic acid (ACC) leads to the inhibition of primary root elongation, the increase of primary root width, and the induction of ectopic root hairs (Masucci and Schiefelbein, 1996; Smalle and Van Der Straeten, 1997; Le et al., 2001). These three ethylene induced responses will promote soil penetration and greater anchorage on the ground.

Great advances in ethylene signaling pathway have been made in the past decade using genetic approaches in Arabidopsis (Merchante et al., 2013). In the absence of ethylene, the receptors and other related proteins recruit the Raf-like kinase CTR1 which phosphorylates the C-terminal end of EIN2, thus preventing it from translocating into the nucleus to stabilize the downstream transcription factors EIN3/EIL1. In the presence of ethylene, the hormone binds to the receptors thus inactivating CTR1, so the unphosphorylated C-terminal end of EIN2 can be cleaved and moves into the nucleus to stabilize EIN3/ EIL1 which will activate the downstream transcriptional cascade (Gao et al., 2003; Ju et al., 2012; Qiao et al., 2012; Wen et al., 2012).

Intriguingly, mutants of auxin synthesis, signaling pathway or transport show aberrant responses to ethylene, indicating crosstalk between these two hormones. For example, mutations in auxin biosynthesis genes such as *ASA1, ASB1, TAR1* and *TAA1* exhibit ethylene-insensitive root phenotypes (Stepanova et al., 2005; Stepanova et al., 2008). *YUC* genes also play key roles in ethylene-mediated root response (Won et al., 2011). Mutants of *AXR2/IAA7* and *AXR3/IAA17* which encode transcription regulators in the auxin signal pathway exhibit insensitive root growth to ethylene (Alonso et al., 2003). PIN2 and AUX1, two of the auxin transport components are also involved in ethylene-mediated root response (Ruzicka et al., 2007).

Plants have a considerable number of transcription factors which play vital roles in the different development process. Among all the families of transcription factors, the HD-Zip family is unique to plant. These proteins display a singular combination of a homeodomain with a leucine zipper working as a dimerization motif. This family consists of 47 members and can be classified into four subfamilies (Ariel et al., 2007). *ATHB1* participates in the determination of leaf cell fate, whereas *ATHB13* and *ATHB23* are involved in cotyledon and leaf development (Aoyama et al., 1995; Nakamura et al., 2006). *HAT2* overexpression lines have a representative phenotype of auxin-overproducing mutants indicating a role in auxin-mediated development (Delarue et al., 1998; Sawa et al., 2002). *PHV, PHB*, and *REV* have similar functions during embryogenesis and leaf polarity determination (Prigge and Clark, 2006). *ATHB10, ATML1*, and *PDF2* play important roles in cell fates establishment by regulating cell layer-specific gene expression (Abe et al., 2003). Although some proteins in this family have been studied well in the past few years, others still need further investigation.

In this study, we report an HD-Zip gene *HB52* which is involved in ethylene-mediated primary root elongation. *HB52* knock-down mutants are insensitive to ethylene in primary root elongation while *HB52* overexpression lines have shortened roots similar to ethylene treated plants. Biochemical and genetic assays showed that *HB52* is a direct target of EIN3. *DR5:GUS* in *HB52* mutants showed altered auxin basipetal transport. Further analyses demonstrated that *HB52* could directly regulate *PIN2, WAG1, and WAG2*. Moreover, a clear PIN1 and PIN3 apical polarity in the stele and PIN2 apical polarity in the cortex were observed in the *HB52* overexpression line. Our results indicate that HB52 plays a vital role in the inhibition of ethylene-induced primary root growth in Arabidopsis and acts as the crosstalk node between ethylene and auxin signaling pathways in primary root elongation.

## Results

### Expression pattern and subcellular localization of HB52

To investigate the expression pattern of *HB52*, we detected the transcription level of *HB52* in different tissues of 4-week old plants by quantitative RT–PCR. The strongest expression was observed in roots followed by stem and rosette leaves (Figure 1A). To further confirm this result, we generated *HB52pro:GUS* transgenic lines. Histochemical analysis of the transgenic lines showed that *HB52* promoter-driven GUS reporter was primarily expressed in the root tip and hypocotyl base of 4-day old young seedlings (Figure 1B, C and D). In 10-day old seedlings, GUS staining was mainly observed in roots and petiole of rosette leaves (Figure 1E). In mature plants, GUS staining was only found in roots (Figure 1F).

**Figure 1.**
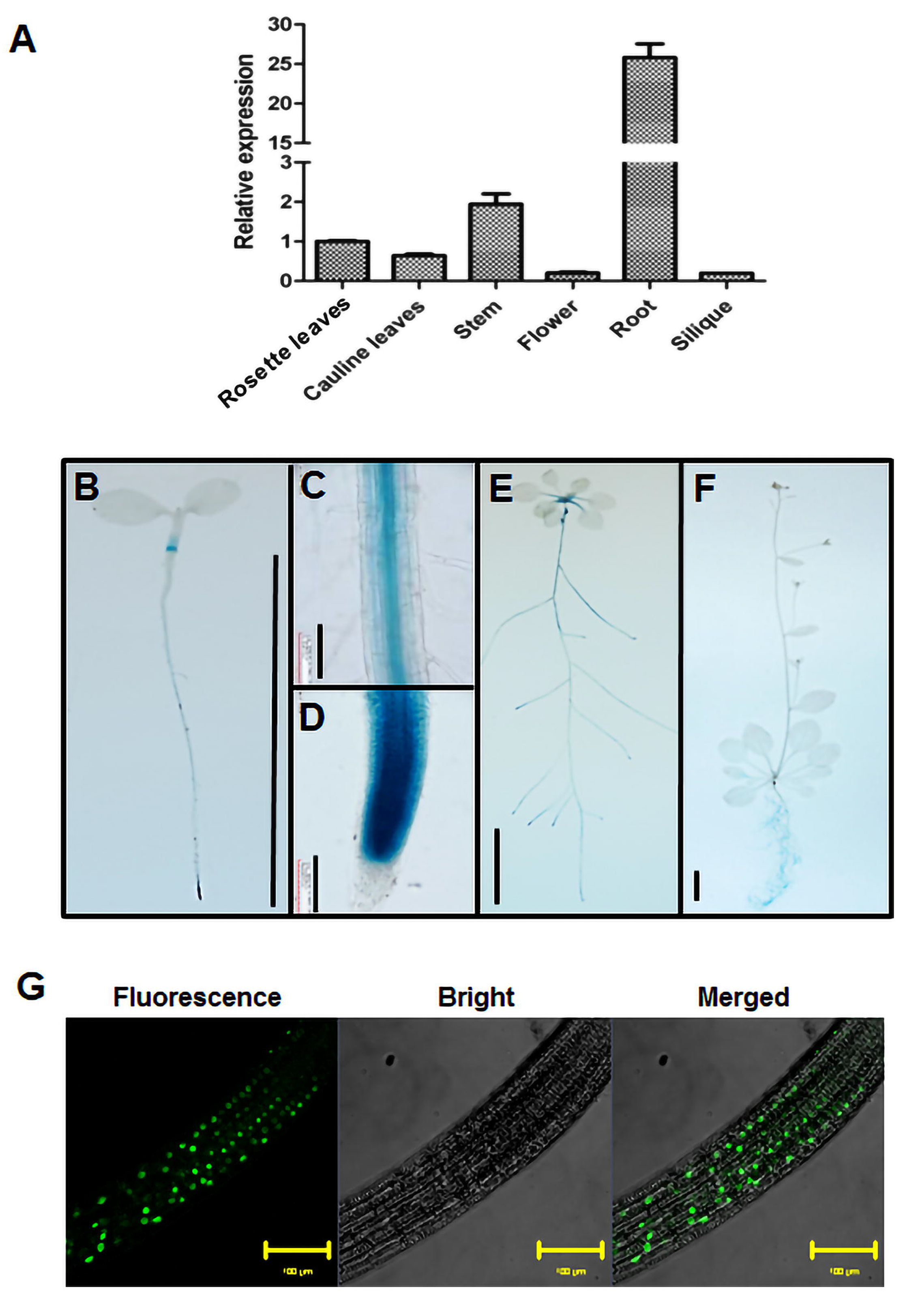
Expression pattern and subcellular localization of HB52. (A) Transcript level of *HB52* in different tissues. Seeds were germinated in the soil for 4 weeks then indicated tissues were collected to isolate RNA and detect the transcript level of *HB52* by quantitative RT–PCR analysis. Values are mean ± SD (n=3 experiments). (B-F) GUS staining of *HB52pro:GUS* transgenic plant. GUS activity was observed in 4-day old seedling (B), 10-day old seedling (E), 4-week adult seedling (F), root of 4-day old seedling (C, D). Plants were incubated in GUS staining solution for 2 hours before photographs were taken. Bar=1cm in B, E, and F. Bar=100μm in C and D. (G) Subcellular localization of the HB52 protein. *35S:HB52-GFP* transgenic seeds were germinated on MS medium for 4 days then fluorescence was observed under confocal laser scanning microscope.(Bar=100μm).

To investigate the subcellular localization of HB52, we generated *35S:HB52-GFP* transgenic lines. Clear fluorescence was observed in the nucleus under confocal laser scanning microscope (Figure 1G). The nucleus localization of HB52 is in coincidence with its function as a transcription factor.

### *HB52* is responsive to ethylene, which depends on ethylene signaling

To confirm whether *HB52* is regulated by ethylene and determine its position in ethylene signaling pathway, we detected the transcript level of *HB52* in the wild type (Col-0) and ethylene signaling mutants using quantitative RT–PCR. *HB52* was upregulated by exogenous ACC (Figure 2A). Moreover, *HB52* was down regulated in ethylene signaling-blocked mutants *ein2-5* and *ein3-1eil1* and upregulated in ethylene signaling-enhanced mutants *35S:EIN3-GFP* and *ctr1-1* without or with exogenous ACC (Figure 2A). To confirm this, we introduced *HB52pro:GUS* into *ein2-5, ein3-1eil1, 35S:EIN3-GFP* and *ctr1-1* background, respectively. The GUS staining of *HB52pro:GUS* was lighter in *ein2-5* and *ein3-1eil1* background while darker in *35S:EIN3-GFP* and *ctr1-1* background when compared with *HB52pro:GUS* without or with exogenous ACC (Figure 2B and 2C). These results indicate that *HB52* acts downstream of *EIN3* and *EIL1*.

**Figure 2.**
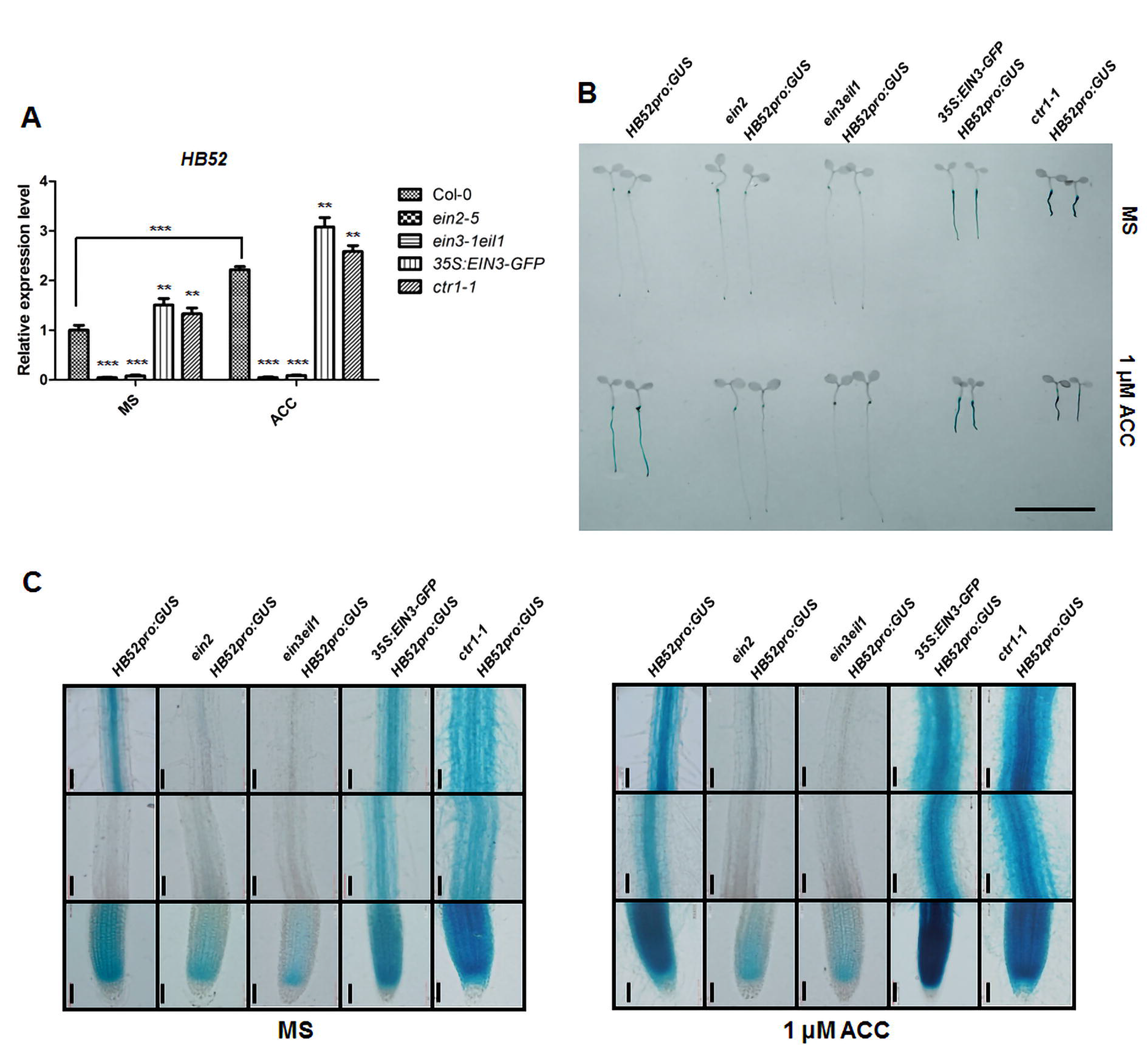
*HB52* is responsive to ethylene and depends on ethylene signaling. (A) Transcript level of *HB52* in Col-0 and ethylene signaling mutants. Seeds were germinated on MS medium for 4 days and then transferred to MS liquid medium without or with 1μM ACC for 24 hours. Then RNA was isolated and quantitative RT–PCR analysis was performed to detect the *HB52* expression level. Values are mean ± SD (n=3 experiments, *P<0.05, **P<0.01, ***P<0.001). Statistically significant differences were calculated based on the Student’s *t*-tests. (B-C) GUS staining of *HB52pro:GUS* transgenic seedlings in ethylene signaling mutants. Seeds were germinated on MS medium for 4 days and then transferred to MS liquid medium without or with 1μM ACC for 24 hours. Seedlings were incubated in GUS staining solution for 0.5 hour before photographs were taken (B). The roots of stained seedlings were observed under a microscope (C). Bar=1cm in B. Bar=100μm in C.

### *HB52* regulates primary root elongation in response to ethylene

To study the role of *HB52* in root elongation in response to ethylene, we obtained the mutant CS909234 with a T-DNA insertion in the promoter of *HB52* from ABRC (Figure S1) and generated an estradiol-induced RNAi line RNAi-6. For clarity, the mutant CS909234 is renamed *hb52*. The transcript level of *HB52* in *hb52* and RNAi-6 was significantly reduced compared with that of the wild type (Figure 3A). Meanwhile, we tried to generate *HB52* overexpression lines driven by 35S promoter but the transgenic plants failed to set seeds due to aberrant development of flowers (data not shown). Therefore, we generated *HB52* overexpression lines driven by estradiol-inducible promoter instead. The transcript level of *HB52* in three representative overexpression lines OX11-5, OX35-2, OX14-1 increased 30, 250 and 870 fold, respectively (Figure 3A). The relative primary root elongation of the three overexpression lines decreased to 87%, 66%, 43% of the wild type after induction (Figure S2B, left panel). Clearly, the primary root length is negatively correlated with *HB52* expression level.

**Figure 3.**
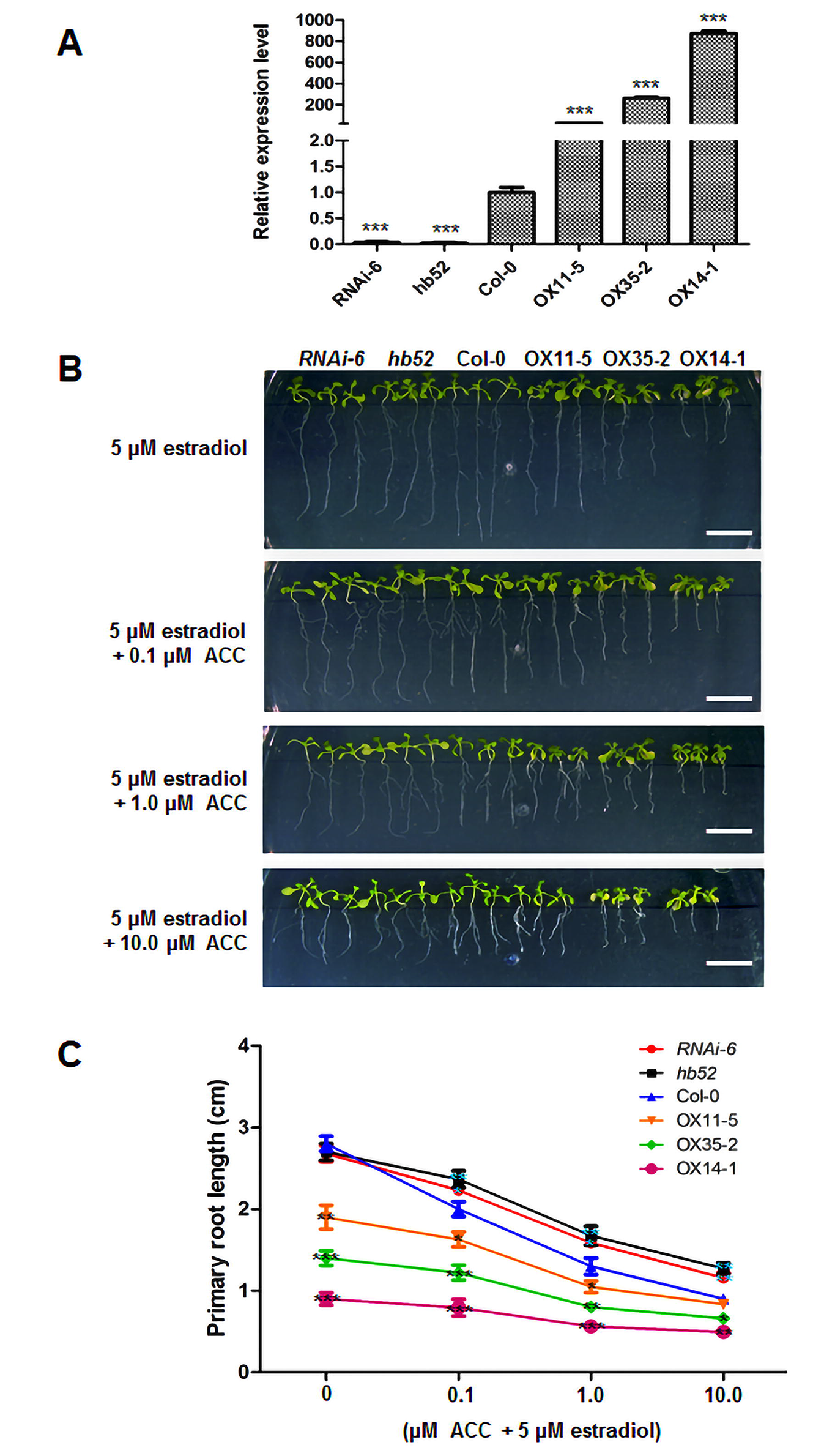
Primary root elongation of *HB52* knock-down mutants and overexpression lines in response to ethylene. (A) *HB52* transcript levels in knock-down mutants and inducible overexpression lines. Seeds were germinated on MS medium for 3 days and the seedlings were then transferred to liquid MS medium with 5μM estradiol for 24 hours to induce gene expression. Then roots were detached and RNA was isolated for quantitative RT–PCR analysis subsequently. Values are mean ± SD (n=3 experiments, *P<0.05, **P<0.01, ***P<0.001). Statistically significant differences were calculated based on the Student’s *t*-tests. (B-C) Root elongation of knock-down mutants and inducible overexpression lines. Seeds were germinated on MS medium for 3 days and then seedlings were transferred to MS medium with 5μM estradiol to induce gene expression for 3 days. Afterwards, seedlings were transferred to MS medium with 5μM estradiol supplemented with 0.1μM ACC, 1μM ACC, and 10μM ACC, respectively for 4 days. Then photographs were taken (B) and primary root length were measured (C). Values are mean ± SD (n=30 seedlings, *P<0.05, **P<0.01, ***P<0.001). Statistically significant differences were calculated based on the Student’s *t*-tests.

If germinated on MS medium with estradiol directly, the overexpression lines OX14-1 exhibited yellow colored cotyledon that might be caused by the high expression level of *HB52* (Figure S2A). To test the response of *HB52* mutants to ethylene and avoid the influence of yellow colored cotyledon, we germinated the seeds on MS medium for 3 days and then transferred the seedlings to MS medium with estradiol for another 3 days to induce gene expression. Afterwards, the seedlings were transferred to MS medium with estradiol supplemented with different concentration of ACC for 4 days to measure primary root elongation. Under 0 μM ACC, the primary root elongation of the knock-down mutants (RNAi-6 and *hb52*) was comparable to that of the wild type control while it was significantly reduced in the overexpression lines, among which the primary root elongation was negatively correlated with *HB52* expression levels (Figure 3B, top panel). In response to ACC, the two *HB52* knock-down lines and three overexpression lines were all less sensitive in root elongation compared with Col-0 (Figure 3B and C). Among the three overexpression lines, OX14-1 is the least sensitive line to ACC in root elongation followed by OX35-2 (Figure 3C). These results indicate that HB52 plays an important role in ethylene-inhibited primary root elongation.

In addition to altered primary root elongation, we observed other root phenotypes associated with varied *HB52* expression levels, which include collapsed root meristem of the overexpression lines (Figure S2A) and altered root gravitropic response of the knock-down mutants and overexpression lines (Figure S3).

### *HB52* is a direct target of EIN3

We have previously shown that *HB52* acts downstream of *EIN3* and *EIL1*. So we next explored whether *HB52* is a direct target of EIN3 and EIL1. Three putative EIN3-binding sites (EBS, TACAT or TTCAAA) were found in the promoter of *HB52* (Konishi and Yanagisawa, 2008; Zhong et al., 2009; An et al., 2012; Li et al., 2013) (Figure 4A). We performed chromatin immunoprecipitation (ChIP) assays using *35S:EIN3-GFP* and *35S:EIL1-GFP* transgenic plants. Marked enrichment of the region containing cis2 site (TACAT) was detected in *35S:EIN3-GFP* transgenic plants by ChIP–PCR assays (Figure 4B and 4C), indicating that EIN3 binds to this region *in vivo*. Furthermore, we conducted yeast-one-hybrid to determine whether EIN3 and EIL1 could directly bind to the EBS in the promoter of *HB52*. The result showed that EIN3 was able to bind to the cis2 site in the promoter of *HB52* (Figure 4D). Taken together, these data suggest that *HB52* is a direct target of EIN3.

**Figure 4.**
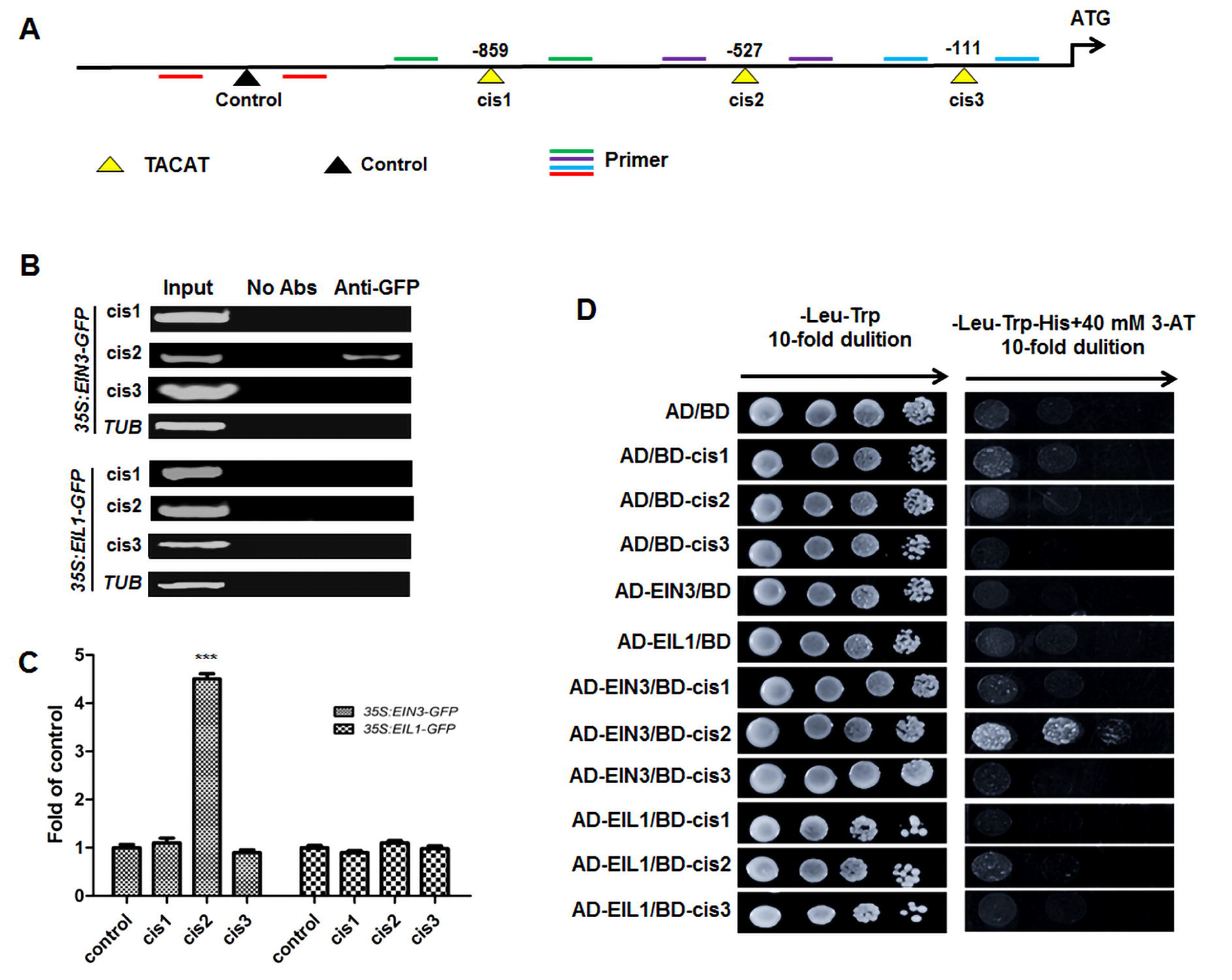
Binding assays of EIN3, EIL1 proteins with the *HB52* promoter. (A) Schematic representation of *HB52* promoter showing putative EIN3 binding sites (EBS) upstream of the transcription start site. EBS are indicated with yellow triangles while black triangle indicates a control that has no EBS in this region. PCR-amplified fragments are indicated by different pairs of colored primers used for ChIP–PCR and quantitative ChIP-PCR. (B-C) ChIP-PCR assays. 4-day old *35S:EIN3-GFP* and *35S:EIL1-GFP* transgenic seedlings were treated with 1μM ACC for 24 hours for ChIP assays. About 200bp *HB52* promoter fragments containing EBS were enriched by anti-GFP antibody in the ChIP-PCR analysis (B). A region of *HB52* promoter which does not contain EBS was used as a control. The results of ChIP–PCR were confirmed by quantitative ChIP–PCR (C). Values are mean ± SD (n=3 experiments, *P<0.05, **P<0.01, ***P<0.001). Statistically significant differences were calculated based on the Student’s *t*-tests. (D) Yeast-one-hybrid assay. pGADT7/EIN3 (AD-EIN3) and pGADT7/EIL1 (AD-EIL1) constructs were co-transformed with pHIS2/HB52 (BD-cis) separately into yeast strain Y187. AD/BD, AD/BD-cis1, AD/BD-cis2, AD/BD-cis3, AD-EIN3/BD, AD-EIL1/BD were used as negative controls.

To further confirm that *HB52* acts downstream of *EIN3*, we crossed *hb52* with *35S:EIN3-GFP* and *ctr1-1* separately*. ctr1-1hb52* had the same point mutation with *ctr1-1* and *35S:EIN3-GFPhb52* had the same expression level of *EIN3* with *35S:EIN3-GFP* (Figure S4A and S4B). *HB52* expression level decreased in *ctr1-1hb52* and *35S:EIN3-GFPhb52* (Figure S4C). The genetic assays showed that the roots of *35S:EIN3-GFPhb52* and *ctr1-1hb52* are longer than that of *35S:EIN3-GFP* and *ctr1-1* without and with exogenous ACC (Figure 5A and 5B). This genetic evidence strongly supports that *HB52* acts downstream of *EIN3*.

**Figure 5.**
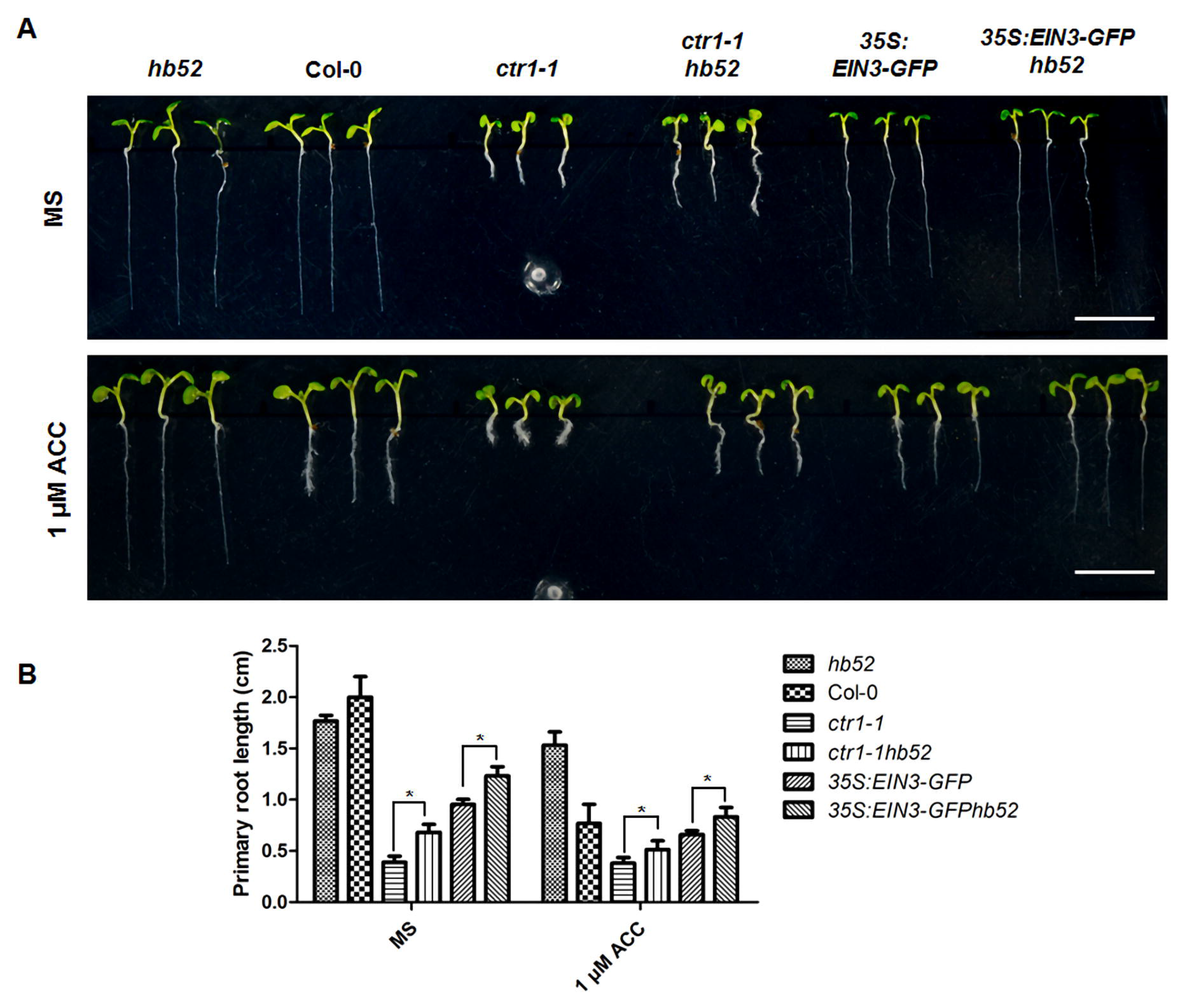
*HB52* genetically acts downstream of *EIN3*. (A) Root elongation phenotype. Seeds of indicated lines were germinated on MS medium without and with 1μM ACC for 5 days before photographs were taken. Bar=1cm. (B) Primary root length. Seeds of indicated lines as in (A) were germinated on MS medium without and with 1μM ACC for 5 days before primary root length was measured. Values are mean ± SD (n=30 seedlings, *P<0.05, **P<0.01, ***P<0.001). Statistically significant differences were calculated based on the Student’s *t*-tests.

### HB52 directly regulates *PIN2, WAG1, and WAG2*

We have noticed that *HB52* knock-down lines and overexpression lines were all insensitive to ACC in root elongation. Obviously, *HB52* plays an important role in ethylene-mediated root elongation. However, the underlying molecular mechanism is unknown. We introduced *DR5-GUS* reporter into *hb52* and OX35-2 background by crossing to see if there is any change of auxin level in root tip. Both lines were confirmed by detecting the transcript level of *HB52* (Figure 6A). Exogenous ACC clearly induces the expression of the *DR5:GUS* reporter in the elongation zone of the wild type but not in the *hb52* background, indicating a blockage in auxin basipetal transport, while the expression of *DR5:GUS* is significantly reduced in the OX35-2 background without or with exogenous ACC (Figure 6B). Taken together, these results suggest that the basipetal transport of auxin is altered by HB52.

**Figure 6.**
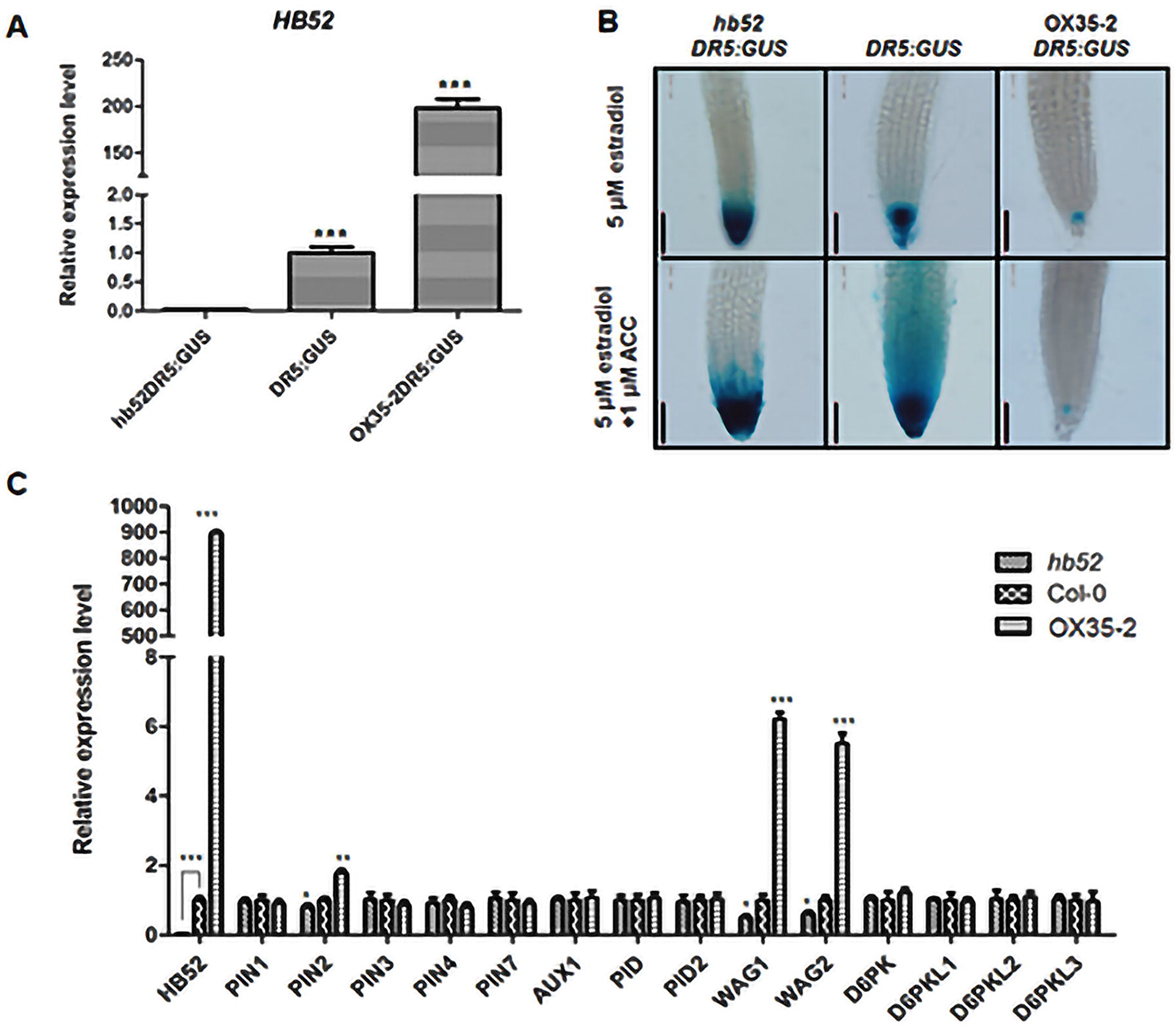
HB52 affects auxin transport by regulating auxin transport-related genes. (A) *HB52* transcript level of *HB52* mutants with DR5:GUS reporter. Seeds of indicated lines were germinated on MS medium for 4 days and transferred to MS liquid medium with 5 μM estradiol for 24 hours. Then roots were detached and RNA was isolated for quantitative RT–PCR analysis subsequently. Values are mean ± SD (n=3 experiments, *P<0.05, **P<0.01, ***P<0.001). Statistically significant differences were calculated based on the Student’s t-tests. (B) GUS staining of DR5:GUS maker lines in varied HB52 backgrounds. Seeds of indicated lines were germinated on MS medium with 5 μM estradiol for 4 days and transferred to liquid MS medium with 5 μM estradiol supplemented without and with 1μM ACC for 24 hours before staining. Seedlings were incubated in GUS staining solution for 2 hours before photographs were taken. Bar=100μm (C) Transcript level of auxin transport-related genes in mutants with varied HB52. Seeds of indicated lines were germinated on MS medium for 4 days and transferred to MS liquid medium with 5 μM estradiol for 24 hours. Then roots were detached and RNA was isolated for quantitative RT–PCR analysis subsequently. Values are mean ± SD (n=3 experiments, *P<0.05, **P<0.01, ***P<0.001). Statistically significant differences were calculated based on the Student’s t-tests.

To investigate the role of HB52 in auxin basipetal transport, we examined the transcript level of *PID, WAG1, WAG2* and other auxin transport related genes. As shown in Figure 6C, *PIN2, WAG1, and WAG2* were downregulated in *hb52* and upregulated in OX14-1. Moreover, several HB52 binding sites were found in promoters of *PIN2, WAG1, and WAG2*, suggesting that these three genes are direct targets of HB52.

To confirm that *PIN2, WAG1, and WAG2* are direct targets of HB52, we demonstrated that HB52 was able to directly bind to at least one homeodomain binding site in the promoter of these three genes by using ChIP-PCR, yeast-one-hybrid, and EMSA (Figure 7, 8 and 9).

**Figure 7.**
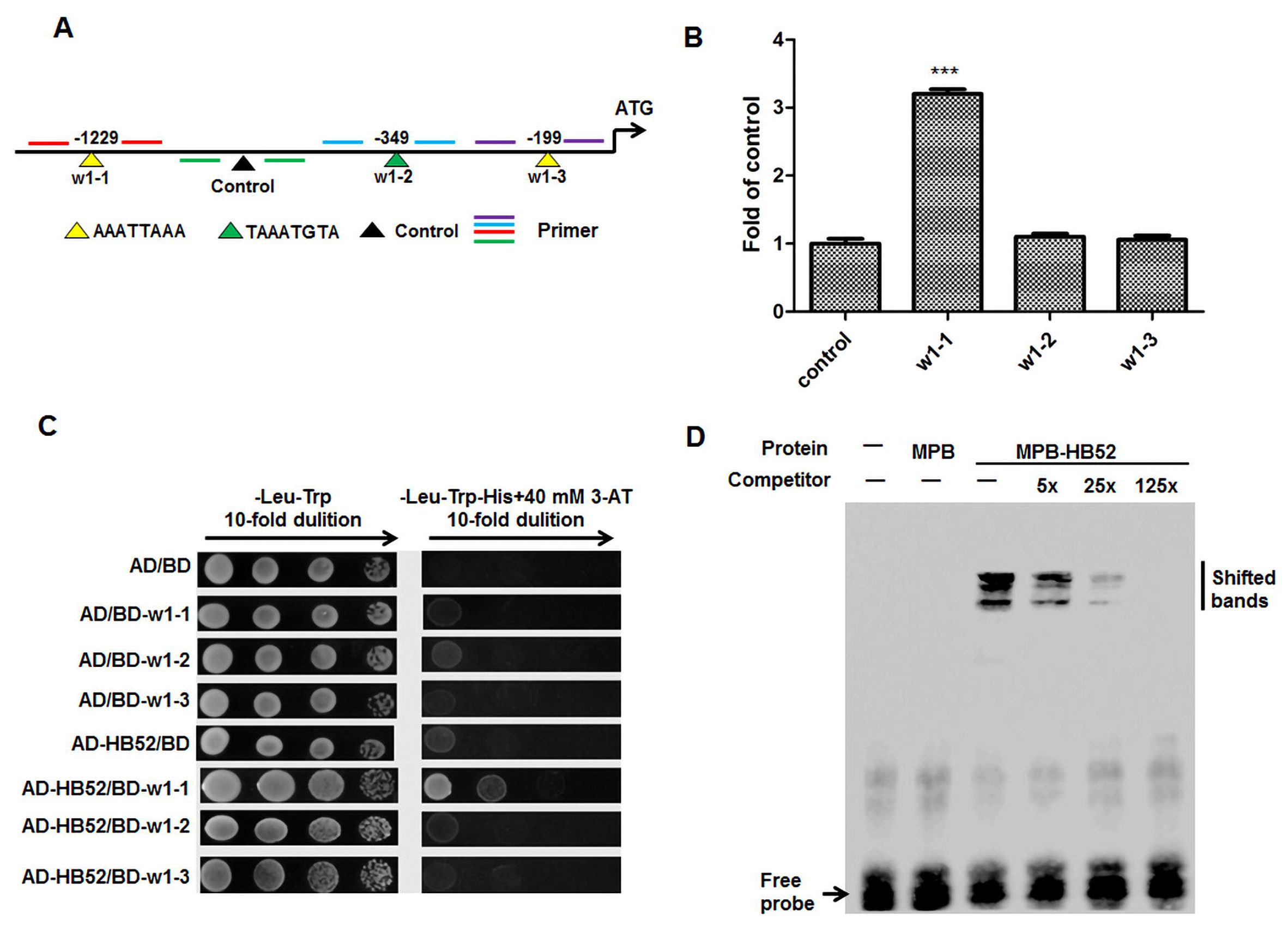
Binding assays of HB52 protein with the *WAG1* promoter. (A) Schematic representation of *WAG1* promoter with putative HB52 binding sites upstream of the transcription start site. HB52 binding sites are indicated with yellow and green triangles while black triangle indicates a control that has no HB52 binding sites in this region. Numbers above the black lines represent the precise HB52 binding sites. PCR-amplified fragments are indicated by different pairs of colored primers and the primers are used to do quantitative RT-PCR. (B) ChIP-PCR assay. 4-day old *35S:HB52-GFP* transgenic seedlings were treated with 1μM ACC for the ChIP-PCR assay. A region of *WAG1* that does not contain HB52 binding sites was used as a control. Values are mean ± SD (n=3 experiments, *P<0.05, **P<0.01, ***P<0.001). Statistically significant differences were calculated based on the Student’s *t*-tests. (C) Yeast-one-hybrid assay. pGADT7/HB52 (AD-HB52) was co-transformed with pHIS2/WAG1 (BD-w1) into yeast strain Y187. AD/BD, AD/BD-w1–1, AD/BD-w1–2, AD/BD-w1–3, AD-HB52/BD were used as negative controls. (D) EMSA of *in vitro* binding. Biotin-labelled probe (w1–1 region) was incubated with HB52-MBP protein. As indicated, HB52-dependent mobility shifts were detected and competed by the unlabeled probe in a dose-dependent manner.

**Figure 8.**
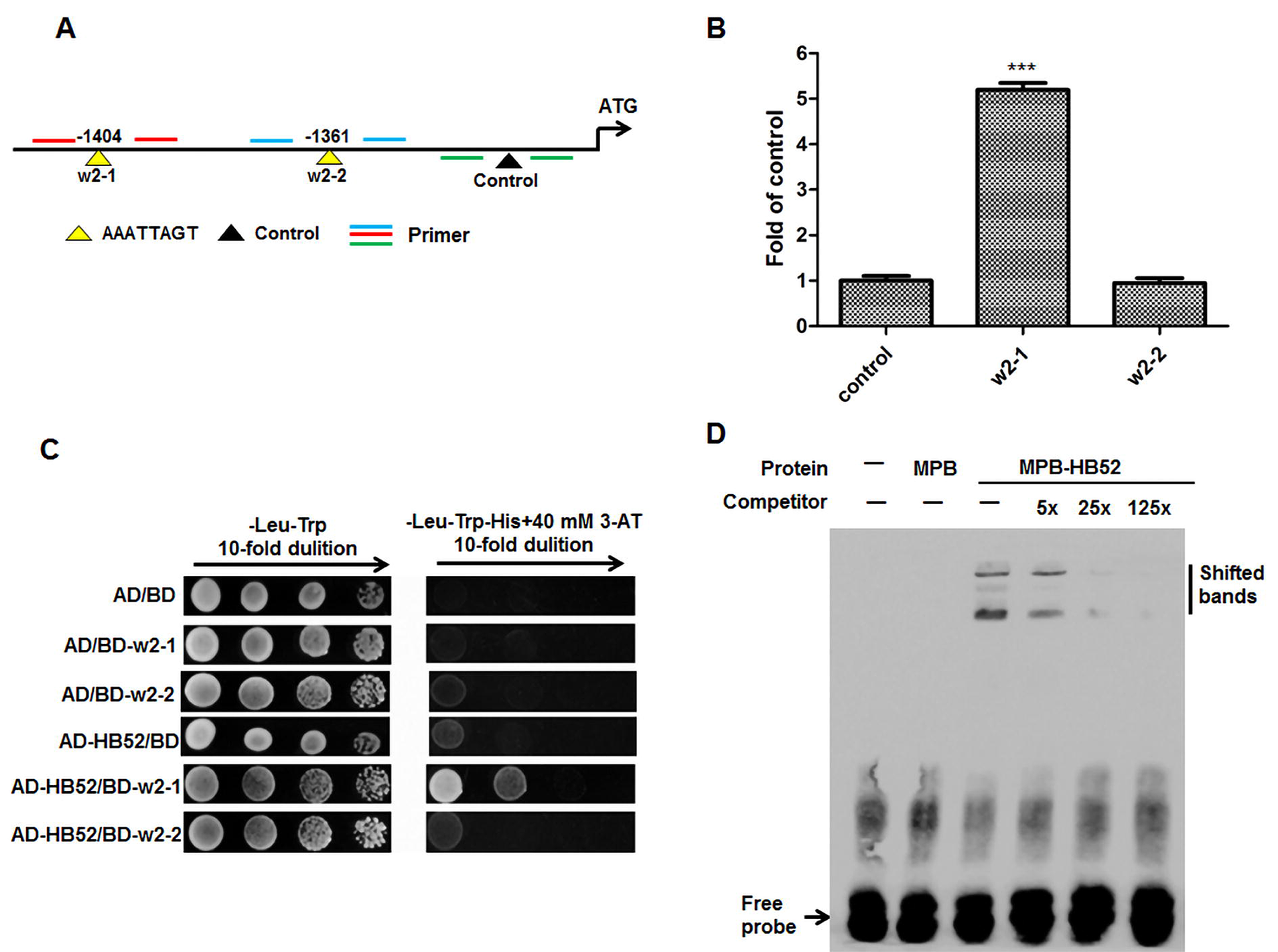
Binding assays of HB52 protein with the *WAG2* promoter. (A) Schematic representation of *WAG2* promoter with putative HB52 binding sites upstream of the transcription start site. HB52 binding sites are indicated with yellow triangles while black triangle indicates a control that has no HB52 binding sites in this region. Numbers above the black lines represent the precise HB52 binding sites. PCR-amplified fragments are indicated by different pairs of colored primers and the primers are used to do quantitative RT-PCR. (B) ChIP-PCR assay. 4-day old *35S:HB52-GFP* transgenic seedlings were treated with 1μM ACC for the ChIP-PCR assay. A region of *WAG2* that does not contain HB52 binding sites was used as a control. Values are mean ± SD (n=3 experiments, *P<0.05, **P<0.01, ***P<0.001). Statistically significant differences were calculated based on the Student’s *t*-tests. (C) Yeast-one-hybrid assay. pGADT7/HB52 (AD-HB52) was co-transformed with pHIS2/WAG2 (BD-w2) into yeast strain Y187. AD/BD, AD/BD-w2–1, AD/BD-w2–2, AD-HB52/BD were used as negative controls. (D) EMSA of *in vitro* binding. Biotin-labelled probe (w2–1 region) was incubated with HB52-MBP protein. As indicated, HB52-dependent mobility shifts were detected and competed by the unlabeled probe in a dose-dependent manner.

**Figure 9.**
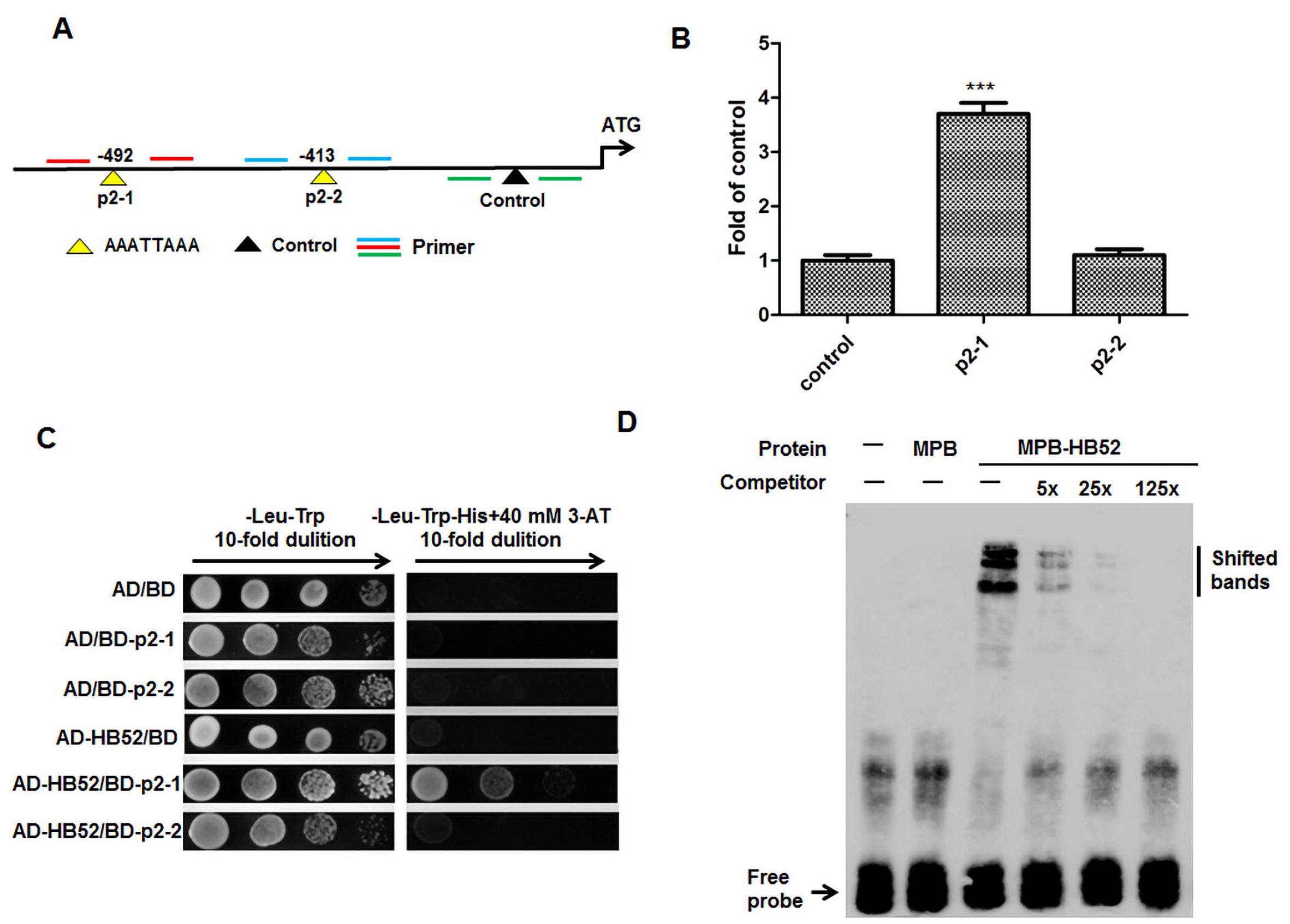
Binding assays of HB52 protein with the *PIN2* promoter. (A) Schematic representation of *PIN2* promoter with putative HB52 binding sites upstream of the transcription start site. HB52 binding sites are indicated with yellow triangles while black triangle indicates a control that has no HB52 binding sites in this region. Numbers above the black lines represent the precise HB52 binding sites. PCR-amplified fragments are indicated by different pairs of colored primers and the primers are used for quantitative ChIP-PCR. (B) ChIP-PCR assay. 4-day old *35S:HB52-GFP* transgenic seedlings were treated with 1μM ACC for the ChIP-PCR assay. A region of *PIN2* that does not contain HB52 binding sites was used as a control. Values are mean ± SD (n=3 experiments, *P<0.05, **P<0.01, ***P<0.001). Statistically significant differences were calculated based on the Student’s *t*-tests. (C) Yeast-one-hybrid assay. pGADT7/HB52 (AD-HB52) was co-transformed with pHIS2/PIN2 (BD-p2) into yeast strain Y187. AD/BD, AD/BD-p2–1, AD/BD-p2–2, AD-HB52/BD were used as negative controls. (D) EMSA of *in vitro* binding. Biotin-labelled probe (p2–1 region) was incubated with HB52-MBP protein. As indicated, HB52-dependent mobility shifts were detected and competed by the unlabeled probe in a dose-dependent manner.

In order to confirm genetically that *PIN2, WAG1*, and *WAG2* act downstream of HB52, we crossed the knockout mutants of *PIN2* (*pin2*, CS8058)*, WAG1* (*wag1*, Salk_002056) and *WAG2* (*wag2*, Salk_070240) with the *HB52* overexpression line (OX35-2), respectively and confirmed the expression of *HB52* (Figure 10A). The results in Figure 10B and 10C show that the primary roots of these hybrid lines are longer than the *HB52* overexpression line with different degrees as predicted. These results suggest that HB52 depends on *WAG1, WAG2*, and *PIN2* for its function in ethylene-mediated root elongation.

**Figure 10.**
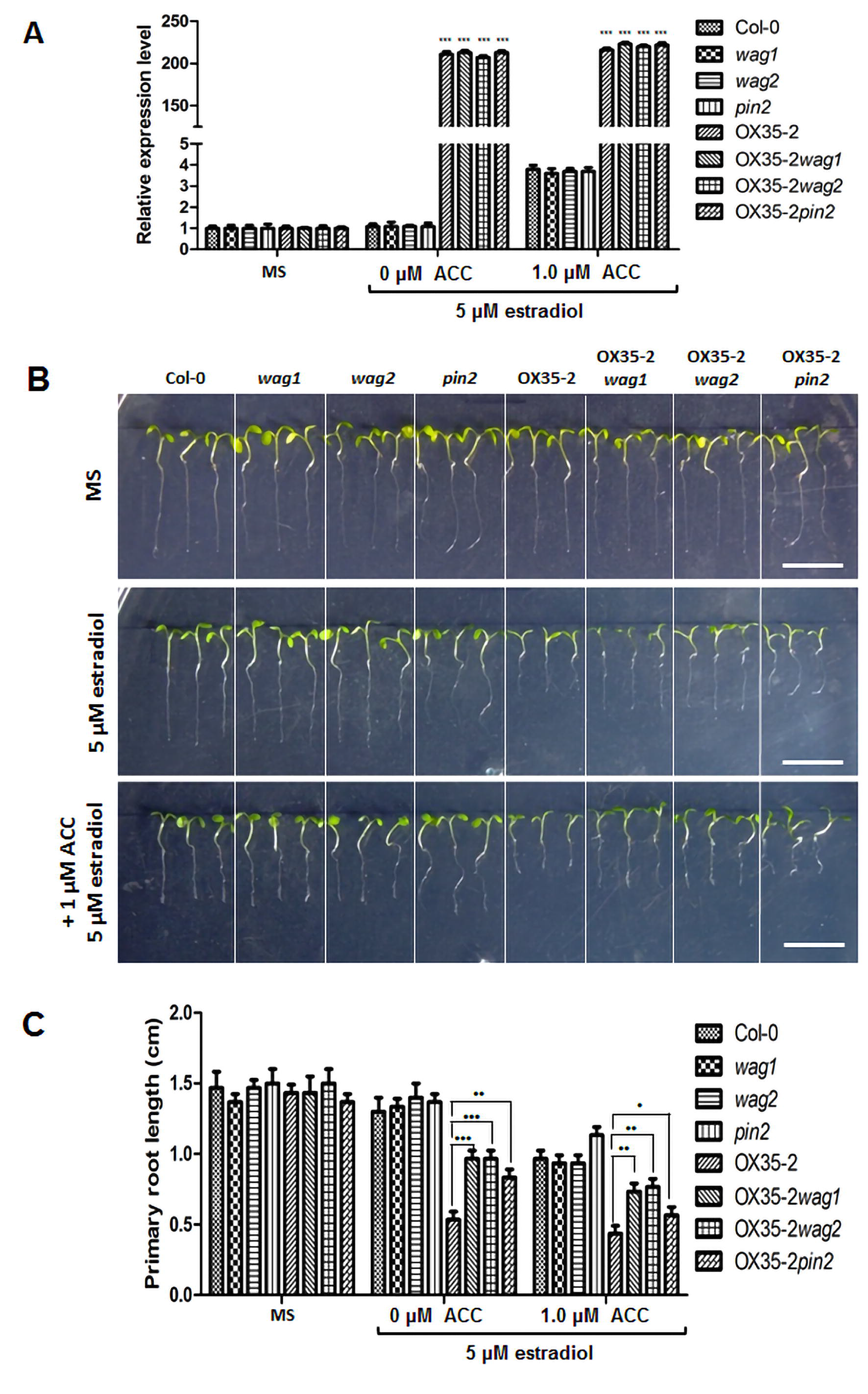
*PIN2, WAG1* and *WAG2* genetically act downstream of *HB52*. (A) HB52 transcript level of varied HB52 mutants. Seeds of indicated lines were germinated on MS medium for 4 days, transferred to MS liquid medium with 5 μM estradiol and MS liquid medium with 5 μM estradiol +1 μM ACC for 48 hours. RNA was isolated for quantitative RT–PCR analysis. Values are mean ± SD (n=3 experiments, *P<0.05, **P<0.01, ***P<0.001). Statistically significant differences were calculated based on the Student’s t-tests. (B-C) Root elongation. Seeds of indicated lines were separately germinated on MS, 5 μM estradiol and 5 μM estradiol +1 μM ACC for 5 days before photographs were taken (B) (Bar=1cm). The primary root length was measured (C). Values are mean ± SD (n=30 seedlings, *P<0.05, **P<0.01, ***P<0.001). Statistically significant differences were calculated based on the Student’s t-tests.

## Discussion

Synergistic effects of auxin and ethylene have been extensively studied in the regulation of root elongation. Ethylene has been shown to increase auxin synthesis, auxin transport to the elongation zone, and auxin signaling at the root tip (Pickett et al., 1990; Alonso et al., 2003; Stepanova et al., 2005; Ruzicka et al., 2007; Swarup et al., 2007; Stepanova et al., 2008; Mao et al., 2016). The HD-Zip transcription factors are a unique family in plants and divided into 4 subfamilies I-IV mainly based on their structure and function. *HB52* belongs to HD-ZIP I and has not been revealed for its role in plants. Members of this subfamily have been shown to be involved in abiotic stress response, ABA-mediated regulation, de-etiolation, and blue-light signaling (Ariel et al., 2007). In this study, we identified that ethylene-responsive *HB52* acts directly downstream of EIN3 to affect auxin basipetal transport by regulating *WAG1, WAG2*, and *PIN2*.

It is known that *HB52* can be upregulated by ethylene in the root in public data such as e-FP browser. A previous study also shows that EIN3, a master regulator of the ethylene signaling pathway, binds directly to the promoter of *HB52* based on the data of EIN3 ChIP-Seq experiments (Chang et al., 2013). So we speculate that it may play a role in ethylene-mediated root regulation. To investigate its function, we first obtained the *HB52* knock-down mutant and overexpression lines. Both *HB52* knock-down mutant and overexpression lines are less sensitive to exogenous ACC in root elongation than wild type (Figure 3). Moreover, the primary roots of *35S:EIN3-GFPhb52* and *ctr1-1 hb52* are longer than *35S:EIN3-GFP* and *ctr1-1* respectively, which further supports the role of *HB52* in ethylene-mediated root elongation (Figure 5). Both ChIP-PCR and yeast-one-hybrid experiments confirm that EIN3 can bind to the promoter of *HB52* (Figure 4), consistent with EIN3 ChIP-Seq data (Chang et al., 2013). The expression pattern of *HB52pro:GUS* reporter in transgenic lines also matches the function of *HB52* in the root (Figure 1 and 2).

To investigate the specific mechanism by which *HB52* controls root elongation. We introduced *DR5:GUS* reporter into *hb52* and OX35-2 background. When treated with ACC, the staining of *DR5:GUS* in *hb52* background showed a blockage in auxin basipetal transport (Figure 6B), which explains the insensitivity of knock-down lines to ethylene (Figure 3B and 3C). The staining of *DR5:GUS* is significantly reduced in OX35-2 background mainly due to the aberrant development of meristematic zone in the root (Figure 6B and S2A). This is the reason why overexpression lines are insensitive to ACC because ethylene-mediated root inhibition needs more auxin basipetal transport from the meristematic zone to the elongation zone (Ruzicka et al., 2007). The aberrant development of meristem is probably the cause of agravitropism (Figure S3) since auxin redistribution in the meristematic zone is of vital importance in regulating gravitropic response (Petrasek and Friml, 2009).

The root phenotype of *HB52* overexpression lines is very similar to that of *PID, WAG1* and *WAG2* overexpression lines. Estradiol-induced overexpression of *PID, WAG1* or *WAG2* led to reduced *DR5:GUS* expression, loss of gravitropism and collapse of root meristem. It was reported that the collapsed root meristem can be rescued by NPA (Benjamins et al., 2001; Dhonukshe et al., 2010). We previously obtained *35S: HB52* lines with severe fertility problems (data not shown) just like the *35S: PID* lines due to abnormal flower development (Benjamins et al., 2001). A frequent collapse of root meristem was observed in overexpression lines and can be rescued by NPA (Figure S2B, right panel). Considering the fact that auxin transport was altered in *HB52* mutants and the overexpression lines had so many similarities with AGC3 kinase overexpression lines (Figure 6B, S2 and S3), we detected the transcript level of the genes related to auxin transport and found *PIN2, WAG1, and WAG2* were downregulated in *HB52* knock-down mutants and upregulated in overexpression lines (Figure 6C).

It has been shown that *pin2/eir1* is insensitive to ethylene in root elongation and exogenous ACC upregulates the *PIN2* expression of *proPIN2:GUS* and *proPIN2:PIN2-GFP*, indicating *PIN2* is involved in the ethylene-mediated root inhibition. But PIN2 is not a direct target of EIN3 (Benjamins et al., 2001; Chang et al., 2013). The link between ethylene and *PIN2* is still to be revealed. *PID, WAG1*, and *WAG2* belong to the plant-specific AGCVIII family of kinases and work redundantly to instruct PIN apical polarity in root development. The most distal cells of the *pidwag1wag2* root epidermis displayed basal localization of PIN2 as compared with its apical localization in wild type, while overexpression of these three genes leads to apically localized PIN1 in the root stele, PIN2 in the cortex and PIN4 in the root meristem (Dhonukshe et al., 2010). It has been demonstrated that PIN2 in the epidermis is responsible for auxin basipetal transport and required for root gravitropic response (Ruzicka et al., 2007). The root of *HB52* knock-down mutant is agravitropic and show partly blocked auxin basipetal transport (Figure S3 and 6B) mainly due to the less apical localization of PIN2 in the epidermis caused by downregulation of *PIN2, WAG1, and WAG2* (Figure 6C). By using yeast-one-hybrid, ChIP-PCR, EMSA, and genetic analyses, we further proved that *PIN2, WAG1, and WAG2* are direct targets of HB52 in ethylene-mediated root inhibition (Figure 7, 8, 9 and 10).

Taken together, our results support a model where ethylene stabilizes EIN3 and upregulates *HB52*. HB52 then increases the expression of *PIN2, WAG1, and WAG2*. As a result, more auxin is transported to the elongation zone, leading to inhibition of root elongation.

## Materials and Methods

### Plant materials and growth conditions

Arabidopsis thaliana ecotype Columbia-0 (Col-0) was used as wild-type. A homozygous *HB52* knock-down mutant CS909234 was ordered from Arabidopsis Biological Resource Center. The OX11-5, 35-2, 14-1, 18-4, RNAi-6, *HB52pro:GUS, 35S:HB52-GFP, 35S:EIN3-GFP* transgenic plants were obtained by Agrobacterium (C58C1) -mediated transformation using the Arabidopsis floral-dip method. For OX11-5, 35-2, 14-1 and 18-4, the *HB52* coding sequence was amplified by pER8-HB52-P1 and pER8-HB52-P2 and cloned into pER8. For RNAi-6, about 200bp of the HB52 coding sequence was amplified by RNAi-P1 and RNAi-P2 and then by RNAi-P3 and RNAi-P4, both segments were cloned into phj33, and then shuttled it into the pER8. For *HB52pro:GUS*, the promoter of HB52 were amplified by GUS-HB52-P1 and GUS-HB52-P2 and cloned into pDONR207, and then shuttled it into the pCB308R. For *35S:HB52-GFP*, the HB52 coding sequence without a stop codon were amplified by GFP-HB52-P1 and GFP-HB52-P2 and cloned into pDONR207, and then shuttled it into the pGWB5.

Several plant materials were previously described: *ein2-5* (Alonso et al., 1999), *ein3-1 eil1-1* (Alonso et al., 2003), *ctr1-1* (Kieber et al., 1993), *35S:EIN3-GFP*. *HB52pro:GUSein2-5, HB52pro:GUSein3-1eil1, HB52pro:GUS35S:EIN3-GFP* and *HB52pro:GUSctr1-1* were crossed by *HB52pro:GUS and ein2-5, ein3-1eil1, 35S:EIN3-GFP* and *ctr1-1* separately. *ctr1-1* CS909234 and *35S:EIN3-GFP* CS909234 were crossed by CS909234 with *ctr1-1* and *35S:EIN3-GFP* separately.

Arabidopsis seeds were surface sterilized in 10% bleach for 15 minutes and washed with distilled water for 6 times. Then the seeds were vernalized at 4°C for 3 days and vertically germinated on 1/2MS medium (Murashige and Skoog). If transferred to soil, all plants were grown under long day conditions (16-h light / 8-h dark) at 22–24°C.

### Histochemical GUS staining and fluorescence observation

Histochemical GUS staining of transgenic plants was performed as previously described (Mao et al., 2016). Images were captured using an OLYMPUS IX81 microscope and HiROX (Japan) MX5040RZ.

Fluorescence observation of GFP transgenic plants was imaged using ZEISS710 confocal laser scanning microscope: 543nm for excitation and 620 nm for emission. Fluorescence observation of Propidium iodide (PI) stained transgenic plants. Seedlings were incubated in 10 mg/mL propidium iodide for 3 minutes and washed twice in water. The stained seedlings were imaged using ZEISS710 confocal laser scanning microscope: 488nm for excitation and 510 nm for emission.

### RT-PCR and quantitative RT-PCR analysis

Total RNA was isolated using TRIzol reagent (Invitrogen) and reversed by TransScript RT kit (Invitrogen). Then cDNA was used for RT-PCR and quantitative RT-PCR. For RT–PCR analysis, the PCR products were amplified and examined on 2% agarose gel. Quantitative RT-PCR was performed on StepOne real-time PCR system using SYBR Premix Ex Taq II kit. Genes expression level was normalized by Ubiquitin5 (UBQ5, At3g62250).

### Yeast-one-hybrid assay

Yeast one-hybrid assay was carried out as described previously (Mao et al., 2016). The coding sequence of proteins was cloned into pAD-GAL4-2.1 (AD vector) and the putative protein binding sites were cloned into pHIS2 (BD vector).

### Starch granules staining

Starch granule staining was performed as described previously (Sabatini et al., 1999).

### ChIP assay

ChIP assay was carried out as described previously (Cai et al., 2014).

### EMSA assay

Competitors were commercially synthesized and free probes were synthesized with biotin labelled at the 5’ end. The coding sequence of HB52 was cloned into pMAL-C2 and the HB52-MBP fusion protein was expressed in Rosseta2 strain. EMSA assay was performed using LightShift™ EMSA Optimization and Control Kit (20148×) according to the manufacturer’s instructions.

## Supplemental information

Figure S1. Identification of the T-DNA insertions in CS909234 (*hb52*).

Figure S2. The phenotype of *HB52* overexpression lines.

Figure S3. Root gravitropic response histogram of *HB52* knock-down mutants and overexpression lines.

Figure S4. Identification of *ctr1-1hb52* and *35S:EIN3-GFPhb52*.

Table S1. Primers used in this study (5’- to -3’).

## Author Contributions

C.X. and Z.M. designed the experiments. Z.M., P.X., J.M., L.Y., Y.Y., and H.T. performed the experiments and data analyses. Z.M. wrote the manuscript. C.X supervised the project and revised the manuscript.

## Acknowledgements

This study was supported by grants from NNSFC (grant no.91417306, 30830075), MOST (2012CB114304). The funders had no role in study design, data collection and analysis, decision to publish, or preparation of the manuscript. We thank ABRC for providing the mutant seeds.

## References

Abe, M., Katsumata, H., Komeda, Y., and Takahashi, T. (2003). Regulation of shoot epidermal cell differentiation by a pair of homeodomain proteins in Arabidopsis. Development 130, 635–643.

Alonso, J.M., and Ecker, J.R. (1999). EIN2, a Bifunctional Transducer of Ethylene and Stress Responses in Arabidopsis. Science 284, 2148–2152.

Alonso, J.M., Hirayama, T., Roman, G., Nourizadeh, S., and Ecker, J.R. (1999). EIN2, a bifunctional transducer of ethylene and stress responses in Arabidopsis. Science 284, 2148–2152.

Alonso, J.M., Stepanova, A.N., Solano, R., Wisman, E., Ferrari, S., Ausubel, F.M., and Ecker, J.R. (2003). Five components of the ethylene-response pathway identified in a screen for weak ethylene-insensitive mutants in Arabidopsis. Proceedings of the National Academy of Sciences of the United States of America 100, 2992–2997.

An, F., Zhang, X., Zhu, Z., Ji, Y., He, W., Jiang, Z., Li, M., and Guo, H. (2012). Coordinated regulation of apical hook development by gibberellins and ethylene in etiolated Arabidopsis seedlings. Cell research 22, 915–927.

Aoyama, T., Dong, C.H., Wu, Y., Carabelli, M., Sessa, G., Ruberti, I., Morelli, G., and Chua, N.H. (1995). Ectopic expression of the Arabidopsis transcriptional activator Athb-1 alters leaf cell fate in tobacco. The Plant cell 7, 1773–1785.

Ariel, F.D., Manavella, P.A., Dezar, C.A., and Chan, R.L. (2007). The true story of the HD-Zip family. Trends in plant science 12, 419–426.

Benjamins, R., Quint, A., Weijers, D., Hooykaas, P., and Offringa, R. (2001). The PINOID protein kinase regulates organ development in Arabidopsis by enhancing polar auxin transport. Development 128, 4057–4067.

Cai, X.T., Xu, P., Zhao, P.X., Liu, R., Yu, L.H., and Xiang, C.B. (2014). Arabidopsis ERF109 mediates cross-talk between jasmonic acid and auxin biosynthesis during lateral root formation. Nat Commun 5, 5833.

Chang, K.N., Zhong, S., Weirauch, M.T., Hon, G., Pelizzola, M., Li, H., Huang, S.S., Schmitz, R.J., Urich, M.A., Kuo, D., Nery, J.R., Qiao, H., Yang, A., Jamali, A., Chen, H., Ideker, T., Ren, B., Bar-Joseph, Z., Hughes, T.R., and Ecker, J.R. (2013). Temporal transcriptional response to ethylene gas drives growth hormone cross-regulation in Arabidopsis. eLife 2, e00675.

Chaves, A.L.S., and Mello-Farias, P.C. (2006). Ethylene and fruit ripening: from illumination gas to the control of gene expression, more than a century of discoveries. Genetics & Molecular Biology 29, 508–515.

Delarue, M., Prinsen, E., Onckelen, H.V., Caboche, M., and Bellini, C. (1998). Sur2 mutations of Arabidopsis thaliana define a new locus involved in the control of auxin homeostasis. The Plant journal: for cell and molecular biology 14, 603–611.

Dhonukshe, P., Huang, F., Galvan-Ampudia, C.S., Mahonen, A.P., Kleine-Vehn, J., Xu, J., Quint, A., Prasad, K., Friml, J., Scheres, B., and Offringa, R. (2010). Plasma membrane-bound AGC3 kinases phosphorylate PIN auxin carriers at TPRXS(N/S) motifs to direct apical PIN recycling. Development 137, 3245–3255.

Gao, Z., Chen, Y.F., Randlett, M.D., Zhao, X.C., Findell, J.L., Kieber, J.J., and Schaller, G.E. (2003). Localization of the Raf-like kinase CTR1 to the endoplasmic reticulum of Arabidopsis through participation in ethylene receptor signaling complexes. The Journal of biological chemistry 278, 34725–34732.

Grbić, V., and Bleecker, A.B. (2003). Ethylene regulates the timing of leaf senescence in Arabidopsis. Plant Journal 8, 595–602.

Ju, C., Yoon, G.M., Shemansky, J.M., Lin, D.Y., Ying, Z.I., Chang, J., Garrett, W.M., Kessenbrock, M., Groth, G., Tucker, M.L., Cooper, B., Kieber, J.J., and Chang, C. (2012). CTR1 phosphorylates the central regulator EIN2 to control ethylene hormone signaling from the ER membrane to the nucleus in Arabidopsis. Proceedings of the National Academy of Sciences of the United States of America 109, 19486–19491.

Kieber, J.J., Rothenberg, M., Roman, G., Feldmann, K.A., and Ecker, J.R. (1993). CTR1, a negative regulator of the ethylene response pathway in Arabidopsis, encodes a member of the raf family of protein kinases. Cell 72, 427–441.

Konishi, M., and Yanagisawa, S. (2008). Ethylene signaling in Arabidopsis involves feedback regulation via the elaborate control of EBF2 expression by EIN3. The Plant journal: for cell and molecular biology 55, 821–831.

Le, J., Vandenbussche, F., Van Der Straeten, D., and Verbelen, J.P. (2001). In the early response of Arabidopsis roots to ethylene, cell elongation is up-and down-regulated and uncoupled from differentiation. Plant physiology 125, 519–522.

Li, Z., Peng, J., Wen, X., and Guo, H. (2013). Ethylene-insensitive3 is a senescence-associated gene that accelerates age-dependent leaf senescence by directly repressing miR164 transcription in Arabidopsis. The Plant cell 25, 3311–3328.

Mao, J.L., Miao, Z.Q., Wang, Z., Yu, L.H., Cai, X.T., and Xiang, C.B. (2016). Arabidopsis ERF1 Mediates Cross-Talk between Ethylene and Auxin Biosynthesis during Primary Root Elongation by Regulating ASA1 Expression. PLoS Genet 12, e1005760.

Masucci, J.D., and Schiefelbein, J.W. (1996). Hormones act downstream of TTG and GL2 to promote root hair outgrowth during epidermis development in the Arabidopsis root. The Plant cell 8, 1505–1517.

Merchante, C., Alonso, J.M., and Stepanova, A.N. (2013). Ethylene signaling: simple ligand, complex regulation. Current opinion in plant biology 16, 554–560.

Nakamura, M., Katsumata, H., Abe, M., Yabe, N., Komeda, Y., Yamamoto, K.T., and Takahashi, T. (2006). Characterization of the class IV homeodomain-Leucine Zipper gene family in Arabidopsis. Plant physiology 141, 1363–1375.

Petrasek, J., and Friml, J. (2009). Auxin transport routes in plant development. Development 136, 2675–2688.

Pickett, F.B., Wilson, A.K., and Estelle, M. (1990). The aux1 Mutation of Arabidopsis Confers Both Auxin and Ethylene Resistance. Plant physiology 94, 1462–1466.

Prigge, M.J., and Clark, S.E. (2006). Evolution of the class III HD-Zip gene family in land plants. Evolution & development 8, 350–361.

Qiao, H., Shen, Z., Huang, S.S., Schmitz, R.J., Urich, M.A., Briggs, S.P., and Ecker, J.R. (2012). Processing and subcellular trafficking of ER-tethered EIN2 control response to ethylene gas. Science 338, 390–393.

Ruzicka, K., Ljung, K., Vanneste, S., Podhorska, R., Beeckman, T., Friml, J., and Benkova, E. (2007). Ethylene regulates root growth through effects on auxin biosynthesis and transport-dependent auxin distribution. The Plant cell 19, 2197–2212.

Sabatini, S., Beis, D., Wolkenfelt, H., Murfett, J., Guilfoyle, T., Malamy, J., Benfey, P., Leyser, O., Bechtold, N., Weisbeek, P., and Scheres, B. (1999). An auxin-dependent distal organizer of pattern and polarity in the Arabidopsis root. Cell 99, 463–472.

Sawa, S., Ohgishi, M., Goda, H., Higuchi, K., Shimada, Y., Yoshida, S., and Koshiba, T. (2002). The HAT2 gene, a member of the HD-Zip gene family, isolated as an auxin inducible gene by DNA microarray screening, affects auxin response in Arabidopsis. The Plant journal: for cell and molecular biology 32, 1011–1022.

Smalle, J., and Van Der Straeten, D. (1997). Ethylene and vegetative development. Physiologia plantarum 100, 593–605.

Stepanova, A.N., Hoyt, J.M., Hamilton, A.A., and Alonso, J.M. (2005). A Link between ethylene and auxin uncovered by the characterization of two root-specific ethylene-insensitive mutants in Arabidopsis. The Plant cell 17, 2230–2242.

Stepanova, A.N., Robertson-Hoyt, J., Yun, J., Benavente, L.M., Xie, D.Y., Dolezal, K., Schlereth, A., Jurgens, G., and Alonso, J.M. (2008). TAA1-mediated auxin biosynthesis is essential for hormone crosstalk and plant development. Cell 133, 177–191.

Swarup, R., Perry, P., Hagenbeek, D., Van Der Straeten, D., Beemster, G.T., Sandberg, G., Bhalerao, R., Ljung, K., and Bennett, M.J. (2007). Ethylene upregulates auxin biosynthesis in Arabidopsis seedlings to enhance inhibition of root cell elongation. The Plant cell 19, 2186–2196.

Wen, X., Zhang, C., Ji, Y., Zhao, Q., He, W., An, F., Jiang, L., and Guo, H. (2012). Activation of ethylene signaling is mediated by nuclear translocation of the cleaved EIN2 carboxyl terminus. Cell research 22, 1613–1616.

Won, C., Shen, X., Mashiguchi, K., Zheng, Z., Dai, X., Cheng, Y., Kasahara, H., Kamiya, Y., Chory, J., and Zhao, Y. (2011). Conversion of tryptophan to indole-3-acetic acid by TRYPTOPHAN AMINOTRANSFERASES OF ARABIDOPSIS and YUCCAs in Arabidopsis. Proceedings of the National Academy of Sciences of the United States of America 108, 18518–18523.

Zhong, S., Zhao, M., Shi, T., Shi, H., An, F., Zhao, Q., and Guo, H. (2009). EIN3/EIL1 cooperate with PIF1 to prevent photo-oxidation and to promote greening of Arabidopsis seedlings. Proceedings of the National Academy of Sciences of the United States of America 106, 21431–21436.

